# Limitations of the radiotheranostic concept in neuroendocrine tumors due to lineage-dependent somatostatin receptor expression on hematopoietic stem and progenitor cells

**DOI:** 10.1101/2024.10.07.615865

**Authors:** Nghia Nguyen, Yu Min, Jennifer Rivière, Mark van der Garde, Sukhen Ghosh, Laura M. Bartos, Matthias Brendel, Florian Bassermann, Ali Azhdarinia, Wolfgang A. Weber, Katharina S. Götze, Susanne Kossatz

## Abstract

Radiopharmaceutical therapy (RPT) has become an effective treatment option for neuroendocrine tumors (NETs) and castration-resistant prostate cancer and is in clinical development for a growing number of indications. One of the major advantages of theranostic RPT is that the distribution of radiopharmaceuticals in the human body can be imaged, and radiation doses to the patient’s organs can be calculated. However, accurate dosimetry may be fundamentally limited by microscopic heterogeneity of radiopharmaceutical distribution. We developed fluorescent analogs of somatostatin-receptor-subtype 2 (SSTR2) targeting Lutetium-177 labelled radiopharmaceuticals that are clinically used in patients with neuroendocrine tumors (NETs) and studied their uptake by hematopoietic stem and progenitor cells (HSPC). Hematopoietic stem cells (HSCs) and multipotent progenitor cells (MPPs) showed high and specific SSTR2-ligand uptake, which was at similar levels as neuroendocrine tumor cells. Furthermore, they displayed a several-fold higher uptake of SSTR2-antagonists than of SSTR2-agonists. HSPC treatment with a ^177^Lu-labelled antagonist and agonist showed a stronger reduction of HSC proliferation by the antagonist. Due to the scarcity of HSCs and MPPs, their contribution to total bone marrow uptake of SSTR2-ligands is not detectable in imaging-based dosimetry studies. This likely explains why SSTR2-antagonists caused pancytopenia in clinical trials despite safe dosimetry estimates. In conclusion, target expression heterogeneity can lead to underestimation of radiopharmaceutical toxicity and should be considered when designing clinical trials for new radiopharmaceuticals. The implications of our findings go beyond somatostatin receptor-targeted radiopharmaceuticals and suggest more generally that first-in-human studies should not only be guided by radiation dosimetry from imaging studies but should also include careful escalation of the administered therapeutic activity. The MMC technology is modular and can be applied to other peptide or protein-based radiopharmaceuticals to study cellular distribution and potential bone marrow uptake prior to clinical testing.

## Introduction

Interest in radiopharmaceutical therapy (RPT) has grown tremendously since randomized controlled trials demonstrated its efficacy in neuroendocrine tumors (NETs) and castration-resistant prostate cancer^1–3^. One of the major advantages of RPT is that it follows the concept of theranostics: patients are selected for RPT by imaging using the radiopharmaceutical labeled with a diagnostic isotope, usually a positron emitter. Using positron emission tomography (PET), the amount of radioactivity in tumor tissue can be quantified to predict the likelihood of response to RPT. PET can also visualize heterogeneity in radiopharmaceutical uptake at the whole-body level, and patients in whom some metastases no longer accumulate the radiopharmaceutical due to lack of target expression can be spared ineffective therapies. For therapeutic isotopes that also emit gamma photons in addition to cell-damaging beta- or alpha-radiation, the radiation dose to tumor tissue and normal organs can be determined by single-photon computed tomography (SPECT)^2^.

Despite these advantages, it is important to note that the spatial resolution of clinically used PET and SPECT is limited to approximately 0.5 and 1 cm, respectively. This means that the activity concentrations measured by these imaging modalities represent averages over 0.125 mL to 1.0 mL. Heterogeneity in radiotracer uptake below this level cannot be resolved. Consequently, small fractions of cells that express the target for a radiopharmaceutical may receive significantly more radiation than predicted by imaging if they are surrounded by cells that do not express the target. This is of particular concern for radiation-sensitive cells such as hematopoietic cells.

Here, we report that this is not only a theoretical concern, but that it can explain the unexpected hematotoxicity of the somatostatin receptor 2 (SSTR2) antagonist ^177^Lu-DOTA-JR11 (also ^177^Lu-satoreotide tetraxetan). This therapeutic radiopharmaceutical had shown promising preclinical data with significantly higher tumor uptake and improved tumor-to-normal organ ratio compared to the clinically used SSTR2 agonists like ^177^Lu-DOTA-TATE (Lutathera)^4–7^. This is attributed to the properties of SSTR2 antagonists, which do not internalize but recognize a much higher number of binding sites on the cell surface compared to internalizing agonists^8^ (**Figure 1a**). However, in a phase I study, 4 of 7 patients experienced prolonged grade 4 hematotoxicity despite extensive pre- and post-therapeutic dosimetry indicating that the bone marrow radiation dose was within safe limits^9^. A subsequent multicenter phase I/II study using lower injected activities of ^177^Lu-DOTA-JR11 also reported substantially higher hematotoxicity than reported for ^177^Lu-DOTA-TATE^10^.

**Figure 1:**
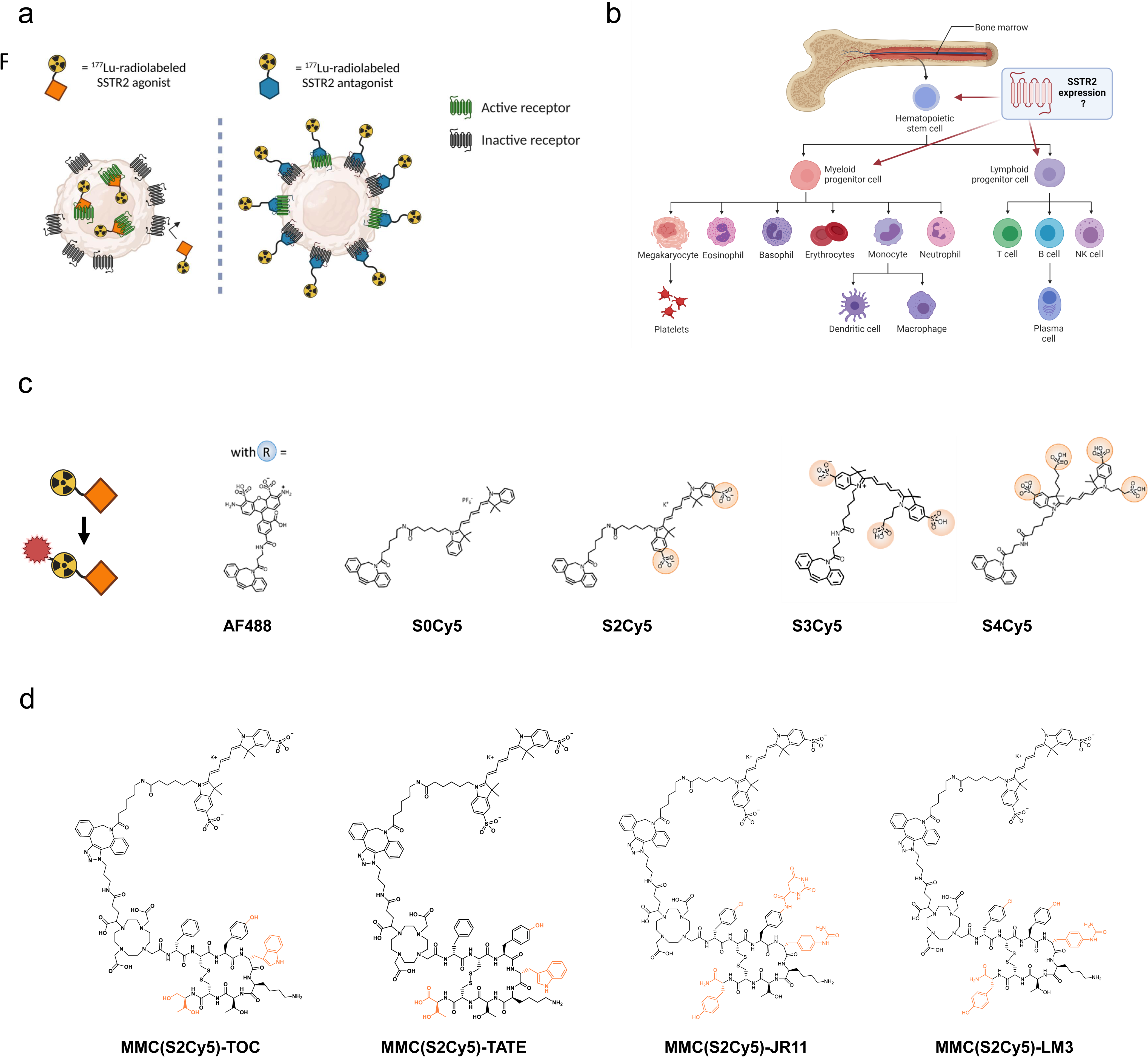
Study concept and outline. **a)** Differences in binding pattern between an agonist and an antagonist to the SSTR2. BioRender.com/k36d510. **b)** Simplified schematic of human hematopoiesis from bone marrow stem cells. BioRender.com/s00l958. **c)** Chemical structures of fluorescent dyes for conjugation onto SSTR2-targeting multimodal peptides. Sulfonic acid groups are circled. **d)** Chemical structures of multimodal agonistic and antagonistic ligands based on the lead dye “S2Cy5”. Highlighted in orange are the chemical differences between the SSTR2-targeting peptides. SSTR2=somatostatin-receptor-2.

This increased hematotoxicity could be the result of SSTR2-expression and hence, specific radioligand binding, on a subpopulation of bone marrow hematopoietic stem and progenitor cells, as we and others have hypothesized^9^. Because of the hierarchical structure of hematopoiesis, this subpopulation may be so small that it has no measurable impact on total bone marrow uptake of radioactivity (**Figure 1b**). To test this hypothesis and identify the mechanism for the higher hematotoxicity of ^177^Lu-DOTA-JR11, we designed fluorescent DOTA-JR11 analogs using a multimodal chelator (MMC)^11^ as linker unit that structurally resembles DOTA and enables bioorthogonal conjugation of the fluorophore. This design allowed us to use radiolabeled versions of the multimodal analogs to ensure they displayed comparable cell-binding characteristics to the clinically used radiopharmaceuticals, and then study their uptake by bone marrow stem cells using fluorescence-activated cell sorting (FACS). In addition to multimodal analogs of ^177^Lu-DOTA-TOC (phase III clinical evaluation; NCT03049189, NCT04919226) and ^177^Lu-DOTA-JR11 (phase I/II clinical evaluation; NCT02592707^10^, NCT04997317, ACTRN12623000185662), we also synthesized multimodal analogs of an additional agonist (^177^Lu-DOTA-TATE, clinically approved for NET treatment)^12^ and antagonist (^177^Lu-DOTA-LM3, first-in-human evaluation)^13^. Subsequently, we analyzed binding to bone marrow hematopoietic stem and progenitor (HSPC) subpopulations and investigated the influence of radioligand treatment on cell viability within these subpopulations.

## Results

### Fluorescent multimodal analogs as suitable tools for cell-based investigations of radioligands

We successfully synthesized a series of multimodal analogs of SSTR2-targeting agonists (based on the clinically used peptides TOC and TATE) and antagonists (based on the ligands JR11 and LM3 in clinical development), using 5 different fluorophores: variants of Cy5 with 0 (S0Cy5), 2 (S2Cy5), 3 (S3Cy5) or 4 (S4Cy5) sulfonic acid groups and AF488 (**Figure 1c**). We chose these dyes since both Cy5 and AF488 are established flow cytometry dyes. The sulfonic acid groups were added to modulate lipophilicity and minimize non-specific binding. The chemical structures of the four S2Cy5 conjugates can be found in **Figure 1d**. Log*D_7.4_* values of all conjugates showed that S2Cy5, S3Cy5 and S4Cy5 conjugates were still hydrophilic (Log*D_7.4_* values < -2), although slightly less compared to the DOTA-parent compounds (Log*D_7.4_* values < -2.5 to -3.5), while the S0Cy5-conjugates were rather lipophilic (Log*D_7.4_* values ∼0) (**Suppl. Fig. 1)**.

To determine the effects of dye labeling on receptor binding and non-specific uptake, we performed radioligand uptake studies with dual-labeled analogs of TOC and JR11 in HCT116-SSTR2 in the presence and absence of octreotide blocking and compared findings to ^177^Lu-DOTA-TOC and ^177^Lu-DOTA-JR11. All ^177^Lu-MMC(Dye)-TOC compounds showed predominantly specific (i.e., blockable) uptake in HCT116-SSTR2 cells at slightly lower levels than ^177^Lu-DOTA-TOC, except for ^177^Lu-MMC(S0Cy5)-TOC, which resulted in high non-specific cell binding (**Figure 2a**). Similar results were observed with the ^177^Lu-MMC(Dye)-JR11 series, with the S0Cy5-JR11 conjugate showing notable non-specific binding (**Figure 2b**). Overall, JR11-based antagonistic ligands showed ca. 10-fold higher cell binding compared to the TOC-based agonistic ligands (ca. 40% of total added activity (taa) vs. 4% taa), which is in agreement with previous studies with TOC and JR11-based radioligands^5–7^. Minimal binding to SSTR2-negative HCT116-WT cells further demonstrated the low non-specific uptake of either ligand (**Suppl. Fig. 2**).

**Figure 2:**
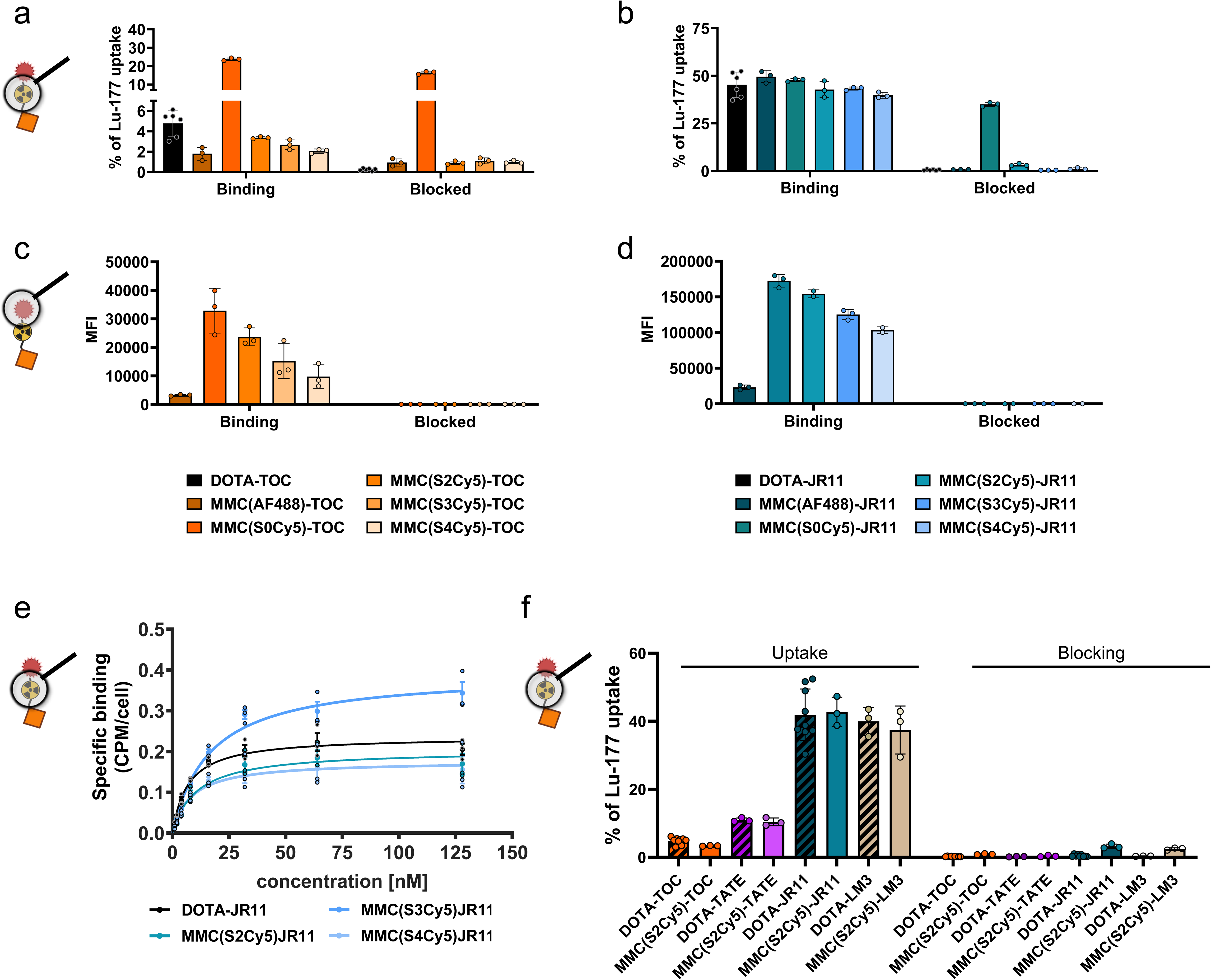
Radioligand and fluorescence brightness characterization on tumor cells. Cellular uptake of ^177^Lu-labeled TOC **(a)** and JR11 **(b)** analogs, reported as % uptake of total added ^177^Lu-activity on HCT116-SSTR2 cells with or without blocking with a 1000-fold excess of DOTA-JR11. Fluorescent readout of cellular uptake of the TOC **(c)** and JR11 **(d)** analogs in HCT116-SSTR2 cells, reported as mean fluorescence intensity with or without blocking with a 100-fold excess of DOTA-JR11. **e)** K_D_ and data represented as saturation binding curve. **f)** Comparison of cellular uptake of all four ^177^Lu-labelled SSTR2-targeting peptides (agonists: TOC, TATE; antagonists: JR11, LM3) reported as % uptake of total added ^177^Lu-activity on HCT116-SSTR2 cells with or without blocking with a 1000-fold excess of DOTA-JR11. Magnifying glass depictions at each figure indicate if the experiment involved the radiolabel or the fluorescent label as readout. SSTR2=somatostatin-receptor-2. MFI=mean fluorescence intensity.

After confirming receptor-mediated binding, we used flow cytometry to determine the brightness of the multimodal conjugates and identify the dye capable of providing the highest detection sensitivity. We found that conjugates with the green-fluorescent dye AF488 yielded much lower mean fluorescence intensities (MFI) compared to the red fluorescent Cy5-series. Due to the resulting lower detection sensitivity, the AF488 conjugates were therefore not further pursued (**Figure 2c, d**). Within the Cy5-series, there was an increasing fluorescence intensity with a decreasing number of sulfonic acid groups across SSTR2 high (HCT116-SSTR2) (**Figure 2c, d**) and low (H69) expressing cell lines (**Suppl. Fig. 3a**). This effect was attributed to the different quantum yields of the dyes, since increasing brightness with a decrease of sulfonic acid groups was also observed using the unconjugated dyes (**Suppl. Fig. 3b**). We also found high binding specificity and strongly increased binding capacity (7-10 fold higher MFI) of the antagonists compared to the agonists in flow cytometry (**Figure 2c, d**), similar to the radioligand binding assay. The S0Cy5-conjugates were abandoned at this point, due to the strong non-specific binding in the radioligand uptake assay. Determination of K_D_-values using HCT116-SSTR2 cells confirmed that no major changes in affinity occurred in the remaining three ^177^Lu-labeled JR11 multimodal analogs compared to ^177^Lu-DOTA-JR11 (**Figure 2e**, **Table 1**). Based on these results, S2Cy5 was selected as the lead dye due to its high brightness. Additional S2Cy5-containing agonist (MMC(S2Cy5)-TATE) and antagonist (MMC(S2Cy5)-LM3 analogs showed similar cell binding and specificity to their clinical counterparts ^177^Lu-DOTA-TATE and ^177^Lu-DOTA-LM3 (**Figure 2f**).

**Table 1:** Determination of KD-values of the multimodal analogs compared to^177^ Lu-DOTA-JR11.

|  | DOTA-JR11 | MMC(S2Cy5)-JR11 | MMC(S3Cy5)-JR11 | MMC(S4Cy5)-JR11 |
| --- | --- | --- | --- | --- |
| $K_D$ (nM) | 6.965 | 10.69 | 16.57 | 8.25 |

Using fluorescence microscopy, all four multimodal ligands exhibited the expected cell-binding properties, evidenced by internalized agonists and cell membrane-bound antagonists (**Figure 3a**).

**Figure 3:**
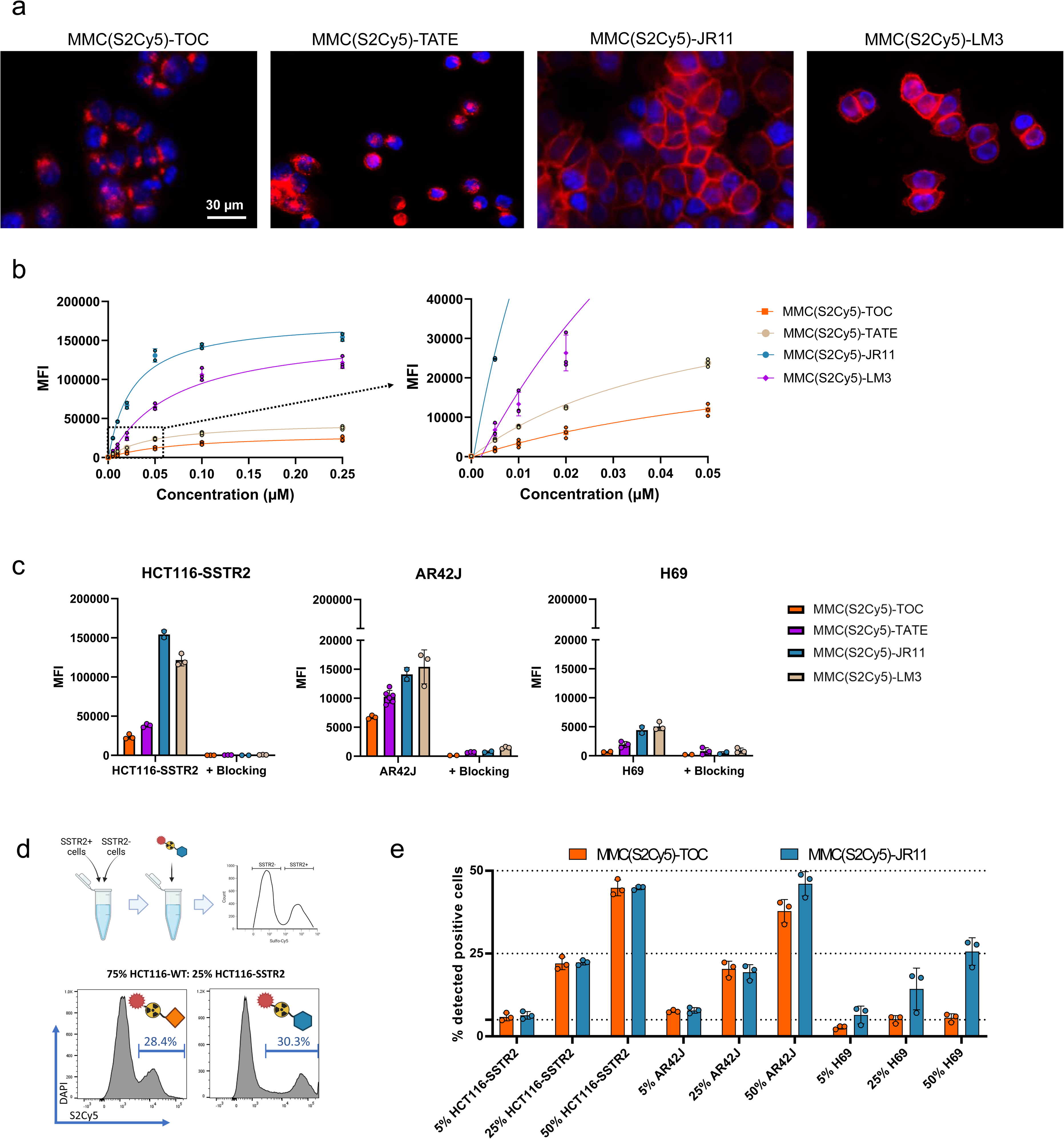
Characterization of MMC(S2Cy5)-ligands on tumor cells. **a)** Fluorescence microscopy to visualize intracellular localisation of MMC(S2Cy5)-ligands in HCT116-SSTR2 cells. S2Cy5 is displayed as red color. Nuclei were stained with Hoechst 33342 (blue). **b)** Concentration dependent cellular uptake of MMC(S2Cy5)-ligands in HCT116-SSTR2 cells. **c)** Cellular uptake of MMC(S2Cy5)-ligands in SSTR2 high (HCT116-SSTR2), medium (AR42J) and low (H69) cells with and without 100-fold blocking with DOTA-JR11. **d)** Schematic depiction of the experiment to determine % positive cells in mixed cell populations. **e)** Mixed cell population experiment using 5, 25 or 50% SSTR2-expressing cells (HCT116-SSTR2, AR42J or H69) mixed with HCT116-WT cells. Data is reported as % of cell that showed binding to MMC(S2Cy5)-TOC/JR11. SSTR2=somatostatin-receptor-2. MFI=mean fluorescence intensity.

Assuming that bone marrow cells might exhibit low levels of SSTR2 expression and hence ligand binding, we aimed to maximize sensitivity by choosing a staining protocol that yields high MFI, while not compromising specificity. Accordingly, we investigated the influence of ligand concentration and incubation time on MFI and were able to detect binding of all ligands using concentrations as low as 5 nM in HCT116-SSTR2 cells, which feature high SSTR2 expression levels (**Figure 3b**). However, higher ligand concentrations showed increased MFI, and hence higher sensitivity. Since we could still achieve complete blocking at 0.25 µM, meaning that no significant nonspecific binding occurred, in high (HCT116-SSTR2), medium (AR42J) and low (H69) expressing cell lines (**Figure 3c**), we chose 0.25 µM for further experiments. We also confirmed that the MFI correlated with the SSTR2-expression levels of the cell line panel (R^2^=0.9627 for MMC(S2Cy5)-JR11 and R^2^=0.9052 for MMC(S2Cy5)-TOC) (**Suppl. Fig. 3c**). The sensitivity limit was reached in the low SSTR2-expressing cell lines H69, where the MFI did not clearly increase above background levels upon incubation with 0.25 µM MMC(S2Cy5)-TOC (**Figure 3c**). However, we observed measurable specific binding in this low-expressing cell line for the other three ligands (MMC(S2Cy5)-TATE/JR11/LM3), confirming high detection sensitivity of these agents. We also conducted further flow cytometric characterization of the binding kinetics and found that the antagonist displayed much faster association and saturation than the agonist (**Suppl. Fig. 3d**). As a final validation step, we prepared mixed cell populations consisting of SSTR2-expressing and non-expressing cells at defined mixing ratios and determined the accuracy of identifying the SSTR2-positive population via binding to the multimodal ligands MMC(S2Cy5)-TOC and -JR11 (**Figure 3d**). In the high (HCT116-SSTR2) and medium (AR42J) expressing cell lines, both the agonist and antagonist were able to quantitatively identify if 5%, 25% or 50% SSTR2-expressing cells were present (**Figure 3e**). In the low-expressing H69 cell lines, both ligands underestimated the % of SSTR2-positive cells, although the antagonist still identified roughly half of the cells, while the agonists did not detect any SSTR2-positive cells. These findings suggest that very low SSTR2-expression levels in bone marrow stem cells could be missed by the multimodal ligands, whereas medium to high SSTR2 expression is detectable.

### SSTR2-targeting peptides show specific binding to hematopoietic stem cells

To investigate binding to bone marrow HSPC, we used human BMMCs and stained them for CD34, CD38, CD90 and CD45RA to identify the following subpopulations: hematopoietic stem cells (HSC, Lin^-^/CD34^+^/CD38^-^/CD45RA^-^/CD90^+^), multipotent progenitors (MPP, Lin^-^/CD34^+^/CD38^-^/CD45RA^-^/CD90^-^), multipotent lymphoid progenitors (MLP, Lin^-^/CD34^+^/CD38^-^/CD45RA^+^/CD90^-^) and committed progenitors (Lin^-^/CD34^+^/CD38^+^) (**Figure 4a**)^14^. Following CD-marker staining, the samples were incubated with 0.25 µM of the respective multimodal ligand and analyzed by flow cytometry. We identified the target subpopulations through standard gating procedures and measured the MFI of the multimodal ligands within each subpopulation (**Suppl. Fig. 4a and 4b** shows examples of gating procedures). During this process, we quantified the frequency of the subpopulations of interest and found that 6.1 ± 2.5% of cells within the entire BMMC population were CD34^+^ (**Figure 4b**). Within the CD34^+^ HSPC population, on average we found 2.7 ± 2.1% HSC, 17.2 ± 9.5% MPP, 6.9 ± 4.4% MLP and 67.0 ±10.8% CD34^+^/CD38^+^ cells **(Figure 4c**).

**Figure 4:**
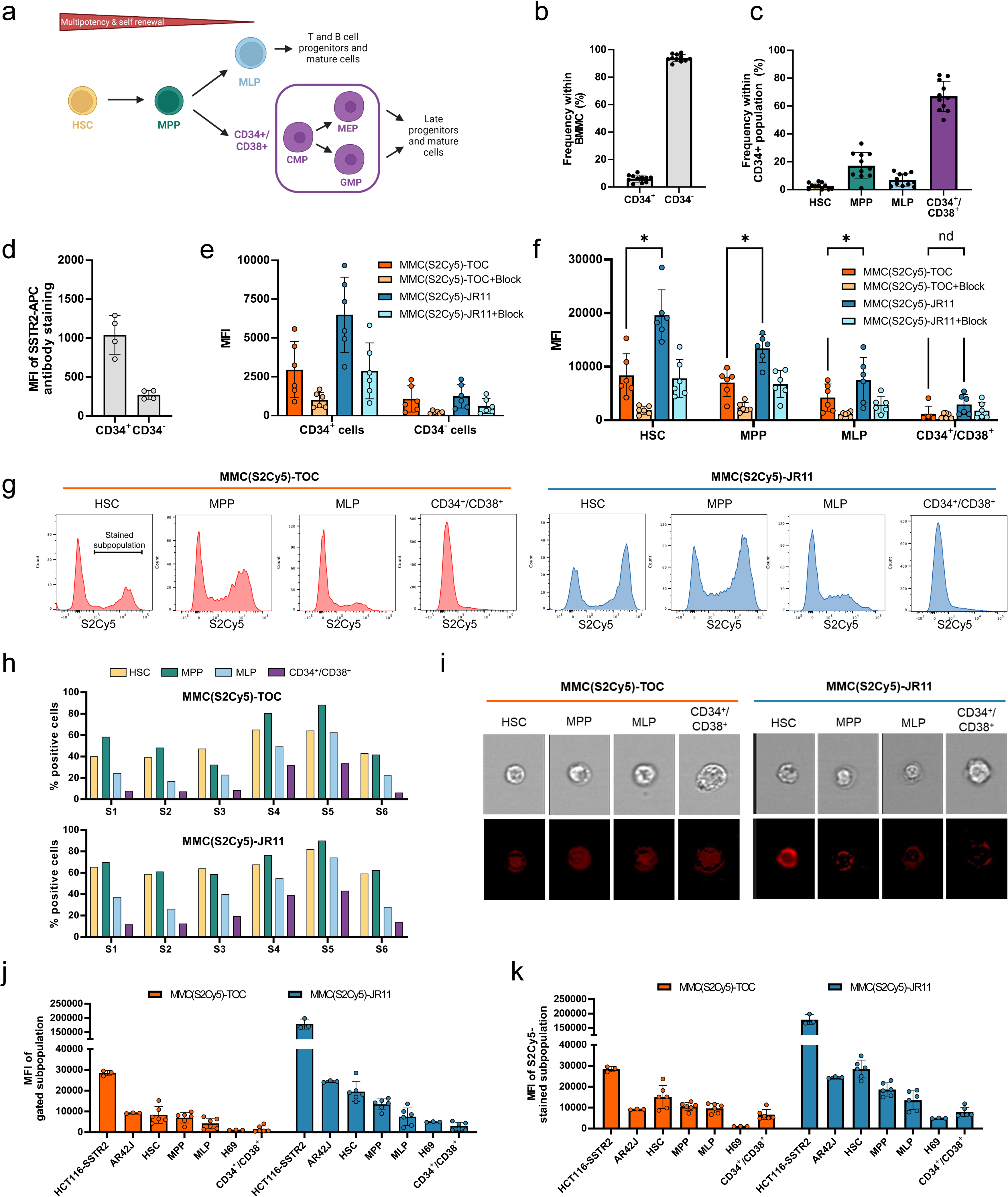
Binding of MMC(S2Cy5)-TOC/JR11 to human hematopoietic stem cells. **a)** Staining strategy to distinguish 4 (HSC, MPP, MLP and CD34+/CD38+) or 6 (HSC, MPP, MLP, CMP, MEP and GMP) human HSPC subpopulations based on CD-markers. BioRender.com/b38j115. **b)** Frequency of CD34+ and CD34-cells within isolated BMMCs. A total of n=6 samples was analyzed in this and subsequent panels. **c)** Frequency of HSC, MPP, MLP-and CD34^+^/CD38^+^ subpopulations within CD34+ HSPC. **d)** SSTR2-antibody binding in CD34+ and CD34-cells. **e)** Binding of MMC(S2Cy5)-TOC/JR11 to CD34+ and CD34-BMMCs with or without blocking with 100-fold excess of DOTA-JR11. **f)** MFI of MMC(S2Cy5)-TOC and -JR11 in HSPC subpopulations with or without blocking. *p<0.01, paired t-test with correction for multiple comparisons. **g)** Histograms of the S2Cy5 signal within HSPC subpopulations showed stained and unstained proportions. **h)** Quantification of the % positive stained cells within each HSPC subpopulation for each analyzed sample. **i)** Imaging flow cytometry to distinguish between internalization or cell-membrane binding of MMC(S2Cy5)-TOC and -JR11 (displayed in red) on HSPC subpopulations. **j)** Comparison of the MFI of known SSTR2-positive tumors cells and HSPC subpopulations after incubation with MMC(S2Cy5)-TOC or -JR11. **k)** Same analysis as in j), but only considering the MFI of the positively stained cells within each subpopulation. SSTR2=somatostatin-receptor-2. MFI=mean fluorescence intensity. HSPC=Hematopoietic stem and progenitor cell.

Staining of HSPC with a fluorescent SSTR2 antibody revealed 4-fold higher binding to CD34^+^ cells compared to CD34^-^ cells (**Figure 4d**). Staining the cells with our multimodal ligands led to similar results, where the uptake of MMC(S2Cy5)-TOC and MMC(S2Cy5)-JR11 was 3 to 5-fold higher in the CD34^+^ population compared to the CD34^-^ population, indicating an SSTR2-mediated binding mechanism (**Figure 4e**). Furthermore, specificity was supported by efficient blocking, which reduced the uptake in all groups. Strikingly, the antagonist MMC(S2Cy5)-JR11 had a 2.2-fold higher uptake than the agonist MMC(S2Cy5)-TOC. Within the subpopulations, the MFI was inversely related to stem cell differentiation status. HSC displayed the highest binding of both compounds (mean MFI: MMC(S2Cy5)-JR11: 19567, MMC(S2Cy5)-TOC: 8352) and a 2.5-fold higher binding of the antagonist compared to the agonist (**Figure 4f**). MPP, MLP and CD34^+^/CD38^+^ showed gradually decreasing MFI. However, we consistently observed a 2 to 3-fold higher uptake of the antagonist compared to the agonist in all subpopulations. We repeated the experiment using the agonist MMC(S2Cy5)-TATE and the antagonist MMC(S2Cy5)-LM3 and found that both agonists (TATE and TOC) as well as both antagonist (LM3 and JR11) showed analogous MFI within the respective subpopulations (**Suppl. Fig. 4c**). Importantly, the more primitive subpopulations showed two distinct peaks in the histograms of HSC, MPP and MLP after incubation with the multimodal TOC/JR11-ligands (**Figure 4g**). This indicates that SSTR2-expressing and non-expressing subpopulations were present within these groups, while the CD34^+^/CD38^+^ committed progenitors showed only one peak with low staining intensity. Quantification of the % positive cells consistently showed that, on average, more than 50% of HSC and MPP stained positive, while it was <40% in MLP and <20% in CD34^+^/CD38^+^ (**Figure 4h**). Using imaging flow cytometry, we confirmed the internalizing property of the agonist and the non-internalizing property of the antagonist on HSPC (**Figure 4i**). We also observed the highest fluorescence intensity of MMC(S2Cy5)-JR11 in HSC, which was in agreement with the flow cytometry MFI analysis.

Subsequently, we wanted to understand if ligand binding is higher or lower than on SSTR2-expressing tumor cells. Therefore, we compared the MFI after incubation of MMC(S2Cy5)-TOC and MMC(S2Cy5)-JR11 within the subpopulations to the known SSTR2-expressing tumor cells HCT116-SSTR2, AR42J and H69. MFI analysis revealed that binding to HSC was only slightly lower than to AR42J cells (**Figure 4i**), which are a widely used model for preclinical investigation of SSTR2-targeted radiopharmaceutical therapy. The lowest MFI were observed in CD34^+^/CD38^+^ cells and in H69 cells, which are a SSTR2-low expressing tumor model. When we compared the MFI only from the positively stained peak of each subpopulation and compared it to the tumor cell lines, we found that the MFI of MMC(S2Cy5)-JR11 was even higher in HSC than in AR42J cells (**Fig. 4j**).

### SSTR2-targeting ^177^Lu-labeled antagonists are more cytotoxic to HSPC than agonists

Finally, we investigated if the differences in JR11/TOC ligand binding in HSPCs also translated into different therapeutic effects of the ^177^Lu-labelled ligands on human HSPC samples *in vitro* (**Suppl. Table 1**). Here, we added CD123 as an additional marker to differentiate between CD34^+^/CD38^+^ subpopulations common myeloid progenitor (CMP; Lin^-^/CD34^+^/CD38^+^/CD45RA^-^/CD123^+^), megakaryocyte-erythrocyte progenitor (MEP; Lin^-^/CD34^+^/CD38^+^/CD45RA^-^/CD123^-^) and granulocyte-monocyte progenitor (GMP; Lin^-^/CD34^+^/CD38^+^/CD45RA^+^/CD123^+^) cells (**Figure 4a**). To assess proliferation within each HSPC subpopulation with or without treatment, cells were labelled with CellTrace Violet prior to the experiment, which allows to measure cell division via dye dilution by flow cytometry (**Figure 5a**). We also assessed the percentage of cell death. Each patient sample was divided into three subsamples (untreated, ^177^Lu-DOTA-TOC, ^177^Lu-DOTA-JR11). For treatment, we added 50 kBq/well of the clinical radioligands ^177^Lu-DOTA-TOC and ^177^Lu-DOTA-JR11 to CTV-stained HSPC in a 6-well plate. Compared to untreated cells, proliferation was reduced across all subpopulations to 77-90 % after ^177^Lu-DOTA-TOC treatment and to 69-79% after ^177^Lu-DOTA-JR11 treatment (**Figure 5b**). However, we noticed a large heterogeneity of the n=16 samples, indicating that certain samples showed a much stronger reaction than others. Therefore, we divided the dataset into “high sensitivity” (n=8 samples with the strongest reduction in proliferation) and “low sensitivity” samples. Within the high-sensitivity samples, the proliferation in HSC was reduced to 82±20% after ^177^Lu-DOTA-TOC treatment and to 42±17% after ^177^Lu-DOTA-JR11 treatment (p=0.001, paired t-test with correction for multiple comparisons). Within all other subpopulations, we also observed lower levels of proliferation in the ^177^Lu-DOTA-JR11 treated samples compared to ^177^Lu-DOTA-TOC, but the differences were not significant. The low-sensitivity cohort showed almost no difference in proliferation between untreated and treated cells. We also analyzed the percentage of dead cells after ^177^Lu-DOTA-TOC/JR11 treatment in the same samples (**Figure 5c**). Compared to untreated cells, cell death was increased in ^177^Lu-DOTA-TOC and ^177^Lu-DOTA-JR11 treated samples. The highest rates of cell death were observed in ^177^Lu-DOTA-JR11 treated samples. The low-sensitivity population showed lower rates of cell death compared to the high-sensitivity cohort. However, variation between the values within each subpopulation was high (up to 10-90% dead cells within one group), and significance levels were not reached.

**Figure 5:**
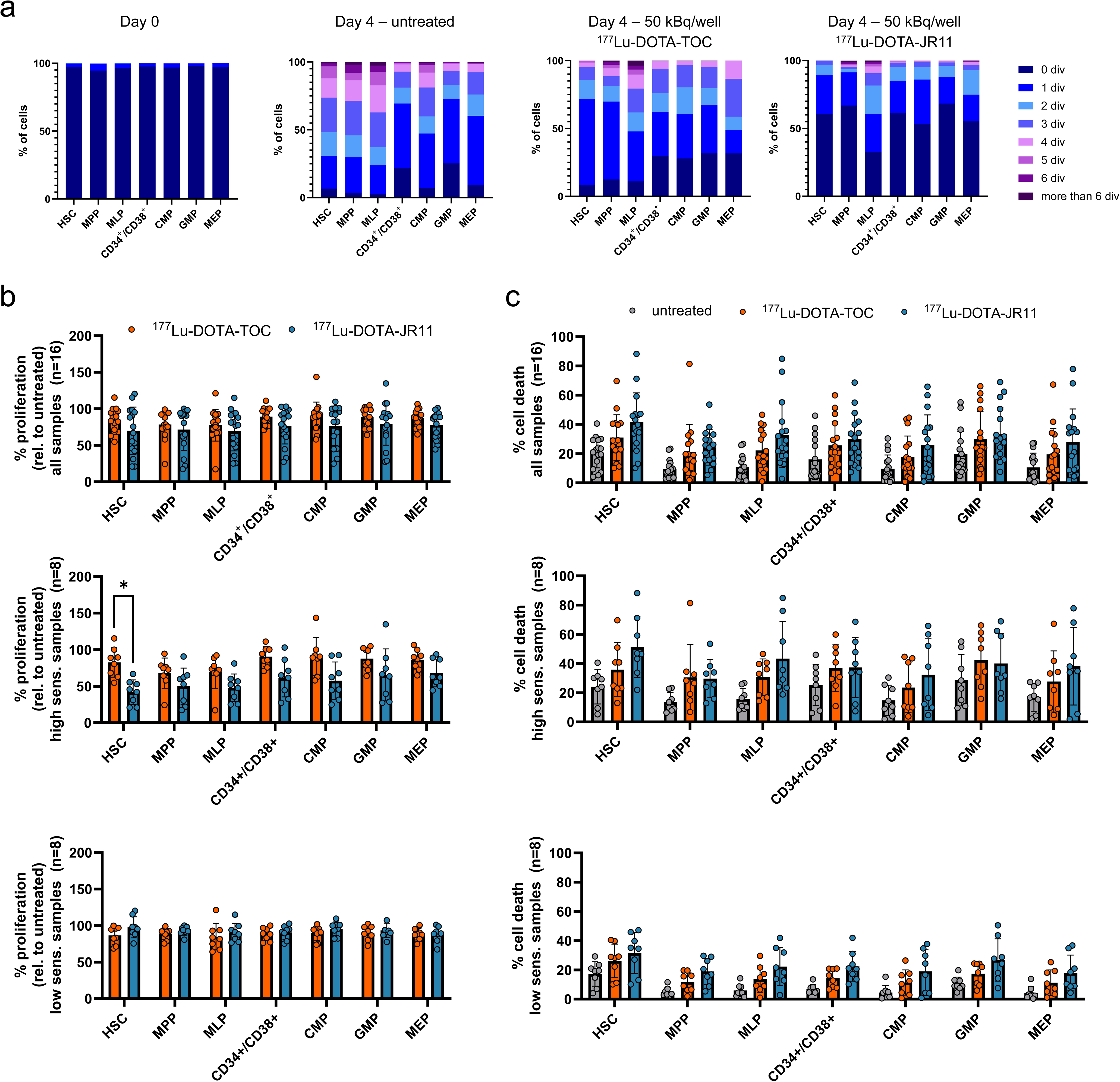
Treatment of HSPC with^177^Lu-DOTA-TOC and -JR11. **a)** Cell proliferation within the HSPC subpopulations was determined by assessing the number of cell divisions with CellTrace™ Violet after 4 days of incubation with and without treatment. Day 0 served as control. **b)** Proliferation score relative to the untreated control within HSPC subpopulations treated with 50 kBq/well of ^177^Lu-DOTA-TOC or ^177^Lu-DOTA-JR11 (n=16). Samples were divided in high and low sensitivity (n=8 each) and analyzed separately. *p=0.001, paired t-test with correction for multiple comparisons. **c)** The same samples were also analyzed for % dead cells in treated and untreated cells. HSPC=Hematopoietic stem and progenitor cell.

## Discussion

In our study, we found that SSTR2-targeting radioligands showed specific binding to subpopulations of bone marrow hematopoietic stem/progenitor cells. Remarkably, the early multipotent and self-renewing HSC and MPP showed similar binding of SSTR2 ligands as neuroendocrine tumor cells. HSC and MPP comprise less than 1% of all bone marrow mononuclear cells and less than 0.005% of the total BM cell population. Hence, radiation dose to these cells cannot be determined by clinical imaging studies, which measure the average radiation dose to the bone marrow. Since HSC are required for the production of all blood cells, severe hematotoxicity may consequently occur at low average bone marrow doses, as was observed in a clinical trial of the new somatostatin receptor antagonist ^177^Lu-DOTA-JR11. Prolonged grade 4 hematotoxicity was observed even though macroscopic bone marrow radiation doses were in a safe range and the total amount of activity administered was at a level that is known to be safe for therapy with somatostatin receptor agonists, such as ^177^Lu-DOTA-TATE^9^. Our data show that SSTR2 antagonists not only had 10-fold higher binding to neuroendocrine tumor cells compared to agonists such as ^177^Lu-DOTA-TOC or -TATE, but also exhibited increased binding to HSC and MPP compared to the agonists.

Oomen et al. reported over 20 years ago on somatostatin receptors in the hematopoietic system^15–17^ and found that SSTR2 is the only expressed SSTR subtype and that it was present in primitive CD34^+^ cells. However, the dose limiting organ for therapy with radiolabeled somatostatin receptor agonists usually is the kidney, and acute hematotoxicity after therapy is rare. Therefore, this observation was not followed up and further subtype analysis into multipotent and progenitor cells has not been performed since then. However, recent clinical experience has shown that the bone marrow has become the dose limiting organ for radioligand therapy with SSTR2 antagonists due to their increased uptake not only into tumor cells, but also into bone marrow cells, which we have shown in this study.

The ability to quantify the distribution of radiopharmaceuticals with imaging studies and calculate radiation doses to target and non-target tissues is one of the major strengths of radiopharmaceuticals, which can allow for remarkably fast clinical development times. For example, ^177^Lu-PSMA-617 went from first synthesis to marketing approval in the US and Europe in 7 years^18,19^. Therefore, there is now enormous interest by academia and industry to develop new therapeutic radiopharmaceuticals^20–22^. However, our data indicate that the development of these agents needs to proceed with caution because microscopic heterogeneity of target expression may result in unexpected toxicity. This does not diminish the value of dosimetry for the development of radiopharmaceuticals in general. Bone marrow dosimetry has been shown to effectively predict toxicity for several agents that bind to more ubiquitously expressed proteins on bone marrow cells (such as CXCR4 ligands)^23,24^ and radiopharmaceuticals without specific binding to bone marrow cells (such as radioiodine)^25,26^. However, in the case of SSTR2-targeted RPT, difficulties in predicting hematologic toxicity based on bone marrow dosimetry have been reported previously^27^.

Our results also demonstrate that multimodal - fluorescent and radioactive - ligands, can be a powerful tool to study the effects of microscopic heterogeneity of target expression. Direct measurement of radioligand binding to bone marrow subpopulations would require the isolation of HSPC subpopulations via cell sorting and subsequent radioligand binding assays but is impractical since the frequencies of the populations of interest are so low that isolation of a high enough number of cells would require prohibitively high sample volumes. Fluorescent dyes, on the other hand, provide a single-cell quantitative readout of binding and do not require subpopulation isolation. However, it is critical to validate that the cellular uptake of the fluorescent ligands closely resembles the uptake by the clinically studied radiopharmaceuticals given the potential for dye labeling to alter SSTR2 binding properties^28^. It is important to note that we did not compare the pharmacodynamic properties, which we assume to be different, but only focused on the cell-binding properties, as those were relevant to our study. Such validations are greatly facilitated by multimodal agents that can also be radiolabeled. We therefore developed new multimodal somatostatin receptor agonists and antagonists using a modified chelator that can be labeled with a radioisotope and a fluorescent dye^29^. Using this approach, we identified fluorescent ligands whose cell-binding properties were analogous to those of clinically used radiolabeled somatostatin receptor ligands. Specifically, the MFI of the fluorescent agents correlated well with the cellular uptake of radioactivity as well as SSTR2-antibody binding. The fluorescent SSTR2 antagonists also showed several-fold higher receptor-specific binding than agonists to tumor cells, as it was the case for the clinically used SSTR2 radioligands. Furthermore, the fluorescently labeled agonists were rapidly internalized, while the antagonists remained at the cell surface. The similar binding patterns on both tumor cells and bone marrow stem cells suggest an SSTR2-mediated binding mechanism in both cases. Using these ligands, we could reliably identify very small fractions of bone marrow hematopoietic cells (< 0.2% of BMMCs) that avidly bound somatostatin receptor ligands.

Importantly, the increased binding of the fluorescent SSTR2 antagonists compared to agonists in HSPC cells resulted in different sensitivity to treatment with SSTR2-targeting radiopharmaceuticals. Specifically, we observed decreased cell proliferation and increased cell death after ^177^Lu-DOTA-JR11 treatment compared to ^177^Lu-DOTA-TOC at the same treatment dose. Thus, our data indicate that the hematotoxicity of SSTR2-targeting radiopharmaceuticals is, to a significant extent, receptor-mediated and that SSTR2 antagonists have a higher potential to induce hematotoxicity than SSTR2 agonists.

However, there are some important considerations when interpreting the data. Due to the very small number of bone marrow cells that overexpress SSTR2, we were unable to directly measure the uptake and retention, and hence radiation dose, of radiolabeled ligands by those cells. However, we expect that the radiation doses to bone marrow cells would be markedly lower than for SSTR2-expressing tumor cells despite comparable radioligand uptake. In solid tumors, a large portion of the dose is generated not by radioligand decay within the tumor cell itself but by the cross-dose from surrounding cells since they are tightly clustered^30,31^. However, HSC and MPP are distributed mostly as single cells or very small clusters in the bone marrow^32^. Hence, it is unlikely that they receive considerable cross-dose from other SSTR2-high expressing cells. In addition, the retention of the SSTR2 antagonists in the HSC may be shorter, because there is less potential for rebinding of ligands that have dissociated from the surface of the cells, which are important mechanisms for their long retention in the tumor tissue^33,34^.

In the radioligand treatment experiment, our data showed relatively large variations of response between samples, leading us to divide the data set into a “sensitive” and “non-sensitive” population, purely based on the reduction of cell proliferation activity. At this point, we do not have any further indicators of what differentiates these samples from one another, since they showed an equal gender and age distribution. However, this biological variability is in line with the clinical observations, where several, but not all patients developed severe hematologic toxicity. It remains unknown at this time if these differences and hematotoxicity were due to a steep dose-response curve where the radiation doses to the bone marrow stem cells were above or below a threshold value in subgroups of patients, or if it reflects differences in radiation sensitivity of bone marrow cells. The variability in response could also be attributed to patient-individual risk factors. HSC and MPP subpopulations are known to have impaired, quiescent or reduced DNA damage repair activity^35,36^. Hence, they could be impacted by the increased irradiation doses more than other, more mature subpopulations. Moreover, HSC aging is associated with a decrease in their regenerative activity, resulting in decreased effector cell production^37^. Any damage to this fragile stem cell subpopulation in older patients receiving RPT would increase possible hematological adverse effects.

Our findings could also be mechanistically related to the occurrence of myelodysplastic syndrome (MDS) or acute myeloid leukemia (AML) following ^177^Lu-DOTA-TATE RPT in NET patients. These therapy-related myeloid neoplasms (t-MN) have been reported in approximately 1-3% of NET patients after different latency periods ranging from several months and many years^38–40^, and even in up to 20% of patients in a heavily pretreated cohort^41^. Different risk factors for developing persistent hematologic dysfunction are currently discussed, including clonal hematopoiesis, chemotherapy with alkylating agents, and short-term hematotoxicity^41–44^. SSTR2 expression on bone marrow stem cells, and hence higher localized radiation doses and DNA damage induction than assumed from dosimetry, could also increase the risk for developing t-MN, especially if paired with another risk factor. Although interesting, our findings should not impact the practice of agonist-based RPT strongly, since t-MN are an established long-term side effect. However, the incidence of t-MN should be carefully observed in future studies employing SSTR2-antagonists since the increased binding to bone marrow stem cells could also increase the risk of developing long-term hematologic dysfunction. So far, the only study reporting long-term side effects of ^177^Lu-DOTA-JR11 reported t-MN in 2/40 patients (5%)^10^.

### Conclusion

Our findings suggest that bone marrow stem and progenitor cell populations, especially HSC and MPP cells, express SSTR2 and bind antagonistic SSTR2-targeting radioligands to a higher extent than agonistic ones, which could explain the increased hematologic toxicity that was clinically observed for SSTR2-targeting antagonists, such as ^177^Lu-DOTA-JR11. The implications of our findings go beyond somatostatin receptor-targeted radiopharmaceuticals and suggest more generally that first-in-human studies of radiopharmaceuticals should not only be guided by radiation dosimetry from imaging studies but should also include careful escalation of the administered therapeutic activity. The MMC technology is modular and can be applied to other peptide or protein-based radiopharmaceuticals to study cellular distribution and potential bone marrow uptake prior to clinical testing. In theranostics, we follow the paradigm that “what we see is what we treat”. Here, we uncovered a case where the treatment unintentionally affected a critical cell population that could not be seen in standard imaging studies, resulting in “sometimes we do not see what we treat”.

## Material and Methods

### Synthesis of multimodal DOTA-JR11/-TOC analogs

The synthesis of multimodal DOTA-JR11/-TOC and DOTA-LM3/-TATE analogs was carried out by replacing the DOTA moiety with an azide-containing cyclen analog, multimodality chelator (“MMC”)^29^, and conjugating DBCO-functionalized dyes using dibenzocyclooctyl (DBCO) copper-free click chemistry (**Figure 1c**). We conjugated 5 dyes: AF488 (Jena Bioscience, Cat. No. CLK-1278-5), Cy5 (“S0Cy5” (Lumiprobe, Cat. No. B30F0), “S2Cy5” (Lumiprobe, Cat. No. 233F0), “S3Cy5” (Jena Bioscience, Cat. No. CLK-A130-5), and “S4Cy5” (Fluoroprobes, Cat. No. 1127-5)) resulting in 5 MMC(Dye)-JR11/-TOC analogs. We synthesized MMC(S2Cy5)-LM3/-TATE variants following the same synthesis strategy. MMC-JR11/-TOC and MMC-LM3/-TATE precursors were purchased from Bachem.

### Preparative RP-HPLC purification

To purify the desired products, reversed-phase high-performance liquid chromatography (RP-HPLC) was performed on an analytical CBM-20A communications bus module, a SPD-M20A prominence diode array detector, a LC-20AP prominence preparative liquid chromatograph with a mobile phase of A = 0.1% TFA or FA in H_2_O, B = 0.1% TFA or FA in MeCN (gradient: 10% - 90% B) at a constant flow rate of 5 mL/min. As a solid phase, a C18-column (Reprosil 120 C18-AQ, 5 μm, 250 x 10 mm, Dr. Maisch, Ammerbuch-Entringen, Germany) was used.

### Analytical RP-HPLC-MS quality control

A Shimadzu RP-HPLC system DGU-20A3 prominence degasser, a LC-30AD Nexera liquid chromatograph (for both pumps) consisting of a SIL-30AC Nexera autosampler, a CBM-20A prominence communications bus module, a RF-20A prominence fluorescence detector, a CTO-20AC prominence column oven, and a SPD-M20A prominence diode array detector) coupled to a Shimadzu liquid chromatograph mass spectrometer LCMS-2020 under usage of a FCV-20AH2 valve unit (Shimadzu) were used to verify the chemical purities of the synthesized compounds.

For the RP-HPLC systems, a mobile phase of A = 0.1% TFA or FA in H_2_O, B = 0.1% TFA or FA in MeCN (gradient: 10% - 90% B) was selected at a constant flow rate of 0.75 mL/min (RP-HPLC coupled to MS). As a solid phase, a C18-column (Reprosil 120 C18-AQ, 5 μm, 150 x 4 mm, Dr. Maisch, Ammerbuch-Entringen, Germany) was used.

#### MMC(AF488)-TOC

AF488-DBCO (0.75 mg, 0.95 µmol) was added to MMC-TOC (1.00 mg, 0.63 µmol) in a mixture of H_2_O and DMSO (1:2.5, 350 µL) and stirred at 37° C for 3 h and then overnight at room temperature (rt). Purification by HPLC with a gradient of 10-90% MeCN (H2O/MeCN + 0.1% TFA) yielded MMC(AF488)-TOC as a blue solid (1.26 mg, 0.53 µmol, 84% yield).

The LC-MS (w/ 10-90% MeCN; H_2_O/MeCN + 0.1% FA) product peak was found at *t*_R_ = 8.6 min. MS, ESI+: m/z calculated 2366.6; found m/z 1185.3 (M+2H^+^), 790.5 (M+3H^+^).

#### MMC(S0Cy5)-TOC

S0Cy5-DBCO (1.00 mg, 1.07 µmol) was added to MMC-TOC (1.00 mg, 0.63 µmol) in DMSO (300 µL) and stirred at 37 °C for 3 h and then overnight at rt. Purification by HPLC with a gradient of 10-90% MeCN (H2O/MeCN + 0.1% TFA) yielded MMC(S0Cy5)-TOC as a blue solid (1.49 mg, 0.63 µmol, 100% yield).

The LC-MS (w/ 10-90% MeCN; H_2_O/MeCN + 0.1% FA) product peak was found at *t*_R_ = 9.9 min. MS, ESI+: m/z calculated 2359.9; found m/z 1180.6 (M+2H^+^), 787.4 (M+3H^+^), 590.9 (M+4H^+^).

#### MMC(S2Cy5)-TOC

S2Cy5-DBCO (1.00 mg, 1.02 µmol) was added to MMC-TOC (1.00 mg, 0.63 µmol) in a mixture of H_2_O and DMSO (3:1, 400 µL) and stirred at 37 °C for 3 h and then overnight at rt. Purification by HPLC with a gradient of 10-90% MeCN (H2O/MeCN + 0.1% TFA) yielded MMC(S2Cy5)-TOC as a blue solid (1.59 mg, 0.63 µmol, 100% yield).

The LC-MS (w/ 10-90% MeCN; H_2_O/MeCN + 0.1% FA) product peak was found at *t*_R_ = 9.0 min. MS, ESI+: m/z calculated 2517.0; found m/z 1260.6 (M+2H^+^), 840.7 (M+3H^+^).

#### MMC(S3Cy5)-TOC

S3Cy5-DBCO (1.00 mg, 0.99 µmol) was added to MMC-TOC (0.75 mg, 0.47 µmol) in H_2_O (820 µL) and stirred at 37 °C for 3 h and then overnight at rt. Purification by HPLC with a gradient of 10-90% MeCN (H2O/MeCN + 0.1% TFA) yielded MMC(S3Cy5)-TOC as a blue solid (0.85 mg, 0.32 µmol, 69% yield).

The LC-MS (w/ 10-90% MeCN; H_2_O/MeCN + 0.1% FA) product peak was found at *t*_R_ = 8.9 min. MS, ESI+: m/z calculated 2585.0; found m/z 1940.2 (3M+4H^+^), 1724.5 (2M+3H^+^), 1293.3 (M+2H^+^), 862.6 (M+3H^+^),

#### MMC(S4Cy5)-TOC

S4Cy5-DBCO (1.00 mg, 0.88 µmol) was added to MMC-TOC (1.00 mg, 0.63 µmol) in H_2_O (400 µL) and stirred at 37 °C for 3 h and then overnight at rt. Purification by HPLC with a gradient of 10-90% MeCN (H2O/MeCN + 0.1% TFA) yielded MMC(S4Cy5)-TOC as a blue solid (0.79 mg, 0.29 µmol, 46% yield).

The LC-MS (w/ 10-90% MeCN; H_2_O/MeCN + 0.1% FA) product peak was found at *t*_R_ = 9.7 min. MS, ESI+: m/z calculated 2707.2; found m/z 1805.9 (2M+3H^+^), 1354.5 (M+2H^+^), 903.4 (M+3H^+^).

#### MMC(AF488)-JR11

AF488-DBCO (0.43 mg, 0.54 µmol) was added to MMC-JR11 (1.00 mg, 0.54 µmol) in a mixture of H_2_O and DMSO (1:1, 800 µL) and stirred at 37° C for 3 h. Purification by HPLC with a gradient of 10-90% MeCN (H_2_O/MeCN + 0.1% TFA) yielded MMC(AF488)-JR11 as an orange solid (0.73 mg, 0.28 µmol, 51% yield).

The LC-MS (w/ 10-90% MeCN; H_2_O/MeCN + 0.1% FA) product peak was found t_R_ = 6.7 min. MS, ESI+: m/z calculated 2634.3: found m/z 1319.3 (M+2H^+^), 880.8 (M+3H^+^).

#### MMC(S0Cy5)-JR11

S0Cy5-DBCO (0.50 mg, 0.54 µmol) was added to MMC-JR11 (0.50 mg, 0.27 µmol) in DMSO (400 µL) and stirred at 37° C for 3 h. Purification by HPLC with a gradient of 10-90% MeCN (H_2_O/MeCN + 0.1% TFA) yielded MMC(S0Cy5)-JR11 as blue solid (0.76 mg, 0.29 µmol, 100% yield).

The LC-MS (w/ 10-90% MeCN; H_2_O/MeCN + 0.1% FA) product peak was found t_R_ = 9.4 min. MS, ESI+: m/z calculated 2626.5: found m/z 1314.1 (M+2H^+^), 876.5 (M+3H^+^), 657.7 (M+4H^+^).

#### MMC(S2Cy5)-JR11

S2Cy5-DBCO (1.00 mg, 1.02 µmol) was added to MMC-JR11 (1.00 mg, 0.54 µmol) in DMSO (1000 µL) and stirred at 37° C for 3 h. Purification by HPLC with a gradient of 10-90% MeCN (H_2_O/MeCN + 0.1% TFA) yielded MMC(S2Cy5)-JR11 as blue solid (0.90 mg, 0.32 µmol, 59% yield).

The LC-MS (w/ 10-90% MeCN; H_2_O/MeCN + 0.1% FA) product peak was found t_R_ = 8.9 min. MS, ESI+: m/z calculated 2784.6: found m/z 1858.9 (2(M+3H^+^), 1394.5 (M+2H^+^), 930.0 (M+3H^+^).

#### MMC(S3Cy5)-JR11

S2Cy5-DBCO (0.79 mg, 0.78 µmol) was added to MMC-JR11 (0.50 mg, 0.27 µmol) in DMSO (1500 µL) and stirred at 37° C for 3 h. Purification by HPLC with a gradient of 10-90% MeCN (H_2_O/MeCN + 0.1% TFA) yielded MMC(S3Cy5)-JR11 as blue solid (0.65 mg, 0.23 µmol, 84% yield).

The LC-MS (w/ 10-90% MeCN; H_2_O/MeCN + 0.1% FA) product peak was found t_R_ = 6.5 min. MS, ESI+: m/z calculated 2852.7: found m/z 1427.9 (M+2H^+^), 952.1 (M+3H^+^).

#### MMC(S4Cy5)-JR11

S2Cy5-DBCO (1.21 mg, 1.07 µmol) was added to MMC-JR11 (1.00 mg, 0.54 µmol) in DMSO (1000 µL) and stirred at 37° C for 3 h. Purification by HPLC with a gradient of 10-90% MeCN (H_2_O/MeCN + 0.1% TFA) yielded MMC(S3Cy5)-JR11 as blue solid (1.19 mg, 0.40 µmol, 74% yield).

The LC-MS (w/ 10-90% MeCN; H_2_O/MeCN + 0.1% FA) product peak was found t_R_ = 9.5 min. MS, ESI+: m/z calculated 2974.8: found m/z 1984.0 (2(M+3H^+^), 1488.3 (M+2H^+^), 992.7 (M+3H^+^).

#### MMC(S2Cy5)-TATE

S2Cy5-DBCO (1.05 mg, 1.07 µmol) was added to MMC-TATE (1.00 mg, 0.63 µmol) in a mixture of H_2_O and DMSO (1:1, 2000 µL) and stirred at 37° C for 3 h and then overnight at rt. Purification by HPLC with a gradient of 10-90% MeCN (H_2_O/MeCN + 0.1% TFA) yielded MMC(S2Cy5)-TATE as blue solid (0.64 mg, 0.25 µmol, 40% yield).

The LC-MS (w/ 10-90% MeCN; H_2_O/MeCN + 0.1% FA) product peak was found t_R_ = 6.9 min. MS, ESI+: m/z calculated 2530.9 (w/o K+): found m/z 845.0 (M+3H^+^).

#### MMC(S2Cy5)-LM3

S2Cy5-DBCO (1.04 mg, 1.06 µmol) was added to MMC-LM3 (1.00 mg, 0.59 µmol) in a mixture of H_2_O and DMSO (1:1, 2000 µL) and stirred at 37° C for 1 h. Purification by HPLC with a gradient of 10-90% MeCN (H_2_O/MeCN + 0.1% TFA) yielded MMC(S2Cy5)-LM3 as blue solid (0.37 mg, 0.14 µmol, 24% yield).

The LC-MS (w/ 10-90% MeCN; H_2_O/MeCN + 0.1% FA) product peak was found t_R_ = 7.3 min. MS, ESI+: m/z calculated 2645.5 (w/o K+): found m/z 883.7 (M+3H^+^).

### 177Lu-radiolabeling

Non-carrier-added ^177^LuCl_3_ was obtained from ITM (Garching, Germany) as an aqueous 0.04 M HCl activity solution. ^177^Lu-radiolabeling of the MMC(Dye) analogs and DOTA-TOC/TATE/LM3/JR11 for cellular uptake, K_D_ assays, and the determination of lipophilicity (log D_7.4_) was carried out in 300 µL dH_2_O (pH 5.3) with 4 nmol of precursor and 20 MBq of ^177^LuCl_3_ at 95 °C for 30 min (molar activity A_m_ = 5 MBq/nmol). Subsequently, 4 nmol of ^nat^LuCl3 was added to saturate the remaining non-labeled chelators. For the treatment of the CD34+ stem cell subpopulations with ^177^Lu-DOTA-JR11/-TOC, the amount of the precursor peptide and ^177^LuCl_3_ were changed to 2 nmol and 400 MBq, respectively (molar activity A_m_ = 200 MBq/nmol). Both reactants were heated to 95 °C for 30 min in 500 µL of 1 M NH_4_OAc buffer (pH = 5.0). Quality control was conducted using radio-TLC and radio-HPLC. For radio-TLCs, we used glass microfiber chromatography paper (ITLC-SG), impregnated with silica gel from Agilent Technologies (Santa Clara, USA) and 0.1 mM citrate buffer. Radio-TLCs were read on an AR-2000 TLC scanner (BIOSCAN).

Radio-RP-HPLC was carried out on a Shimadzu RP-HPLC system, equipped with a a NaI(Tl) well-type scintillation counter from Elysia-Raytest (Straubenhardt, Germany). Radio-HPLC was conducted using a 10-90% MeCN gradient + 0.1% TFA in 15 min on a C18-column (Reprosil 120 C18-AQ, 5 μm, 250 x 10 mm, Dr. Maisch, Ammerbuch-Entringen, Germany), and with 150 kBq activity injection per run. Experiments were only carried out if radio-TLC and radio-HPLC showed >96% radiochemical purity.

### n-Octanol–PBS distribution coefficients (log D_7.4_)

For lipophilicity determination, samples of 0.5 MBq ^177^Lu-labeled compounds in 500 µL PBS were added to 500 µL n-octanol (n = 9), vortexed for 3 min and centrifuged for 5 min at 5000 ×g. 100 µL of each fraction were then measured on a Wizard^2^ automated gamma counter (PerkinElmer), and the logD_7.4_ values were calculated using the formula: counts per minute of octanol phase/counts per minute of water phase.

### Cell culture

H69, AR42J, HCT116-WT (wild type) (all ATCC) and HCT116-SSTR2 (kindly provided by the group of Ali Azhdarinia, UTH Health, Houston, Texas, USA) were cultivated in monolayer culture at 37°C in a 5% CO_2_ humidified atmosphere, following standard procedures. H69 were cultivated as suspension cell line. Cells were maintained in their respective growth medium (Roswell Park Memorial Institute medium containing 10% fetal bovine serum and 1% penicillin/streptavidin for HCT116-WT, HCT116-SSTR2 and H69. The medium for HCT116-SSTR2 cells contained an additional 0.1 mg/mL of Zeocin. Ham’s F-12K containing 20% fetal bovine serum and 1% penicillin/streptavidin for AR42J). Trypsin-EDTA (0.25%) was used to harvest cells and in vitro experiments were carried out at 80% confluency. Cells were authenticated and regularly tested for mycoplasma contamination.

### Cellular uptake of radioligands

SSTR2-dependent uptake of ^177^Lu-MMC(Dye) analogs and reference DOTA-compounds was determined in HCT116-WT and HCT116-SSTR2 cells. Cells were seeded in suspension into a 96-well V-bottom plate at 100,000 cells/well and incubated with the ^177^Lu-labeled MMC(Dye) analogs or ^177^Lu-DOTA-JR11/LM3/TOC/TATE at a final concentration of 6 nM at 37 °C for 60 min. 1000-fold molar excess of DOTA-JR11 was added 10 min prior for blocking. At the end of the incubation time, cells were pelleted at 600×g for 4 min and washed twice with PBS, followed by cell lysis using 1 M NaOH. Cell lysates were measured on a Wizard^2^ automated gamma counter (PerkinElmer). The cellular uptake of the radioligands was quantified as percentage of total radioactivity added. Mean values and standard deviation was calculated from 3-6 biological repeats.

### Determination of the dissociation constant K_D_

K_D_-values of ^177^Lu-MMC(Dye)-JR11 analogs were investigated by incubating HCT116-SSTR2 cells (100,000 cells/well) with increasing concentrations of ^177^Lu-MMC(Dye)-JR11 analogs (1-128 nM) at 37 °C for 60 min. DOTA-JR11 (6 µM) was used to block specific binding at each concentration. At the end of the incubation time, cells were lysed with 1 M NaOH, and measured on a Wizard^2^ automated gamma counter (PerkinElmer). Specific binding was obtained by subtracting non-specific binding (block) from total binding. The obtained counts per minute (specific binding) were then divided by the number of cells (100,000) to obtain the CPM/cell. K_D_ values were obtained by applying a non-linear regression curve fit (Saturation binding – One site specific binding) in GraphPad Prism 10.

### Brightness of multimodal conjugates

The fluorescence intensity of the Cy5-DBCO dyes (“S0Cy5”, “S2Cy5”, “S3Cy5”, and “S4Cy5”) and their JR11/LM3/TOC/TATE-Cy5 dye conjugated equivalents were compared at a final concentration of 5 µM in 150 µL DMSO/well in a flat bottom 96-well plate. Analysis was performed using a Synergy HT MultiMode Microplate Reader (Biotek) at the excitation:emission wavelengths of 590/10 nm:645/20 nm.

### Fluorescence microscopy

Fluorescence microscopy was conducted using an EVOS M7000 fluorescence microscope (Thermo Fisher Scientific) to evaluate the antagonistic and agonistic properties of the MMC(Dye)-JR11/TOC and MMC(S2Cy5)-LM3/TATE. Cells were seeded (30,000 cells/well) on an 8-well chamber slide (Ibidi, #80807) 24 h before the experiment, followed by incubation with 1 µM of MMC(Dye)-JR11/TOC and MMC(S2Cy5)-LM3/TATE at 37 °C for 30 min. Cell nuclei were counterstained with 10 µg/ml Hoechst 33342 (Invitrogen, #H3570). Visualization of the fluorescent compounds was achieved by using the fluorescent filters of GFP (for AF488 dye), and Cy5 (for Cy5 dye).

### Flow cytometric characterization of MMC(Dye)-JR11/TOC analogs

FACS analysis was performed on a BD Canto II (BD Biosciences, USA) equipped with three lasers: blue (488 nm, air-cooled, 20 mW solid-state), red (633 nm, 17 mW HeNe), and violet (405 nm, 30 mW solid-state). DAPI (0.1 µg/mL, D1306, Thermo Scientific) staining was used to exclude dead cells. Single-stained tubes were utilized for multi-color compensation. BD FACSDivaTM software was used for data acquisition and Flowjo 9.0-10.10 (BD Biosciences, USA) was employed for flow cytometric data analysis. To analyze the binding of MMC(Dye)-JR11/TOC, data were analyzed as mean fluorescence intensity (MFI) of the entire population of interest, unless stated otherwise. MMC(S2Cy5)-JR11/TOC was detected in the APC-A channel. To enable comparability, all settings (e.g. compensation, voltage, gain) were standardized within each experiment.

### Dye-dependent signal intensity after cell binding (Sensitivity)

HCT116-SSTR2, HCT116-WT, AR42J or H69 cells were seeded into FACS tubes at 1×10^6^ cells/mL in 1 mL FACS buffer (PBS containing 5% FBS and 0.1% NaN3 sodium azide). The samples were incubated with a final concentration of 0.25 µM MMC(Dye)-TOC/JR11 (Dye = AF488, S0Cy5, S2Cy5, S3Cy5 or S4Cy5) for 30 min at room temperature (rt). For blocking, 25 µM DOTA-JR11 was added 15 min before the multimodal conjugate. The experiment was also carried out for MMC(S2Cy5)-TATE and MMC(S2Cy5)-LM3. Before flow cytometry, samples were washed with 1 mL FACS buffer three times and 100 µL DAPI (0.1 µg/mL) was added as live/dead stain. MFI was calculated from three biological repeats.

### Concentration-dependent signal intensity saturation

HCT116-SSTR2 cells were seeded into FACS tubes at the concentration of 1×10^6^ cells/mL in 1 mL FACS buffer. The samples were incubated with different final concentrations (0.005 µM, 0.01 µM, 0.02 µM, 0.05 µM, 0.1 µM, 0.25 µM) of MMC(S2Cy5)-TOC/JR11 for 30 min at rt. Samples were washed with 1 mL FACS buffer three times and 100 µL DAPI (0.1 µg/mL) was added as live/dead stain prior to the flow cytometry. MFI was calculated from three biological repeats.

### Cell binding kinetics

To determine the binding kinetics of the multimodal analog MMC(S2Cy5)-JR11/TOC via flow cytometry, HCT116-WT, HCT116-SSTR2, AR42J and H69 cells were seeded into FACS tubes at the concentration of 1×10^6^ cells/mL in 1 mL FACS buffer. Cells were incubated with a final concentration of 0.25 µM MMC(S2Cy5)-TOC/JR11 for 2, 5, 15, 30, 60 or 120 min at rt. Samples were washed with 1 mL FACS buffer three times and 100 µL DAPI (0.1 µg/mL) was added as live/dead stain prior to the flow cytometry.

### Identification of SSTR2-positive populations in pre-mixed cultures of SSTR2+/SSTR2-cells

We assessed the ability and sensitivity of MMC(S2Cy5)-JR11/TOC to quantitatively identify SSTR2-positive populations with high, medium or low SSTR2 expression levels in samples of mixed cell populations of known origin (SSTR2+/SSTR2-) in preparation for the bone marrow experiments. HCT116-SSTR2, AR42J and H69 cells were mixed in three proportions with HCT116-WT cells (50%:50%, 25%:75% or 5%:95%) in a concentration of 10^6^ total cells/mL in FACS buffer. Mixed cell cultures were incubated with 0.25 μM MMC(S2Cy5)-JR11/TOC for 30 min at rt. The samples were washed with 1 mL FACS buffer three times and 100 µL DAPI (0.1 µg/mL) was added as live/dead stain prior to flow cytometry. In the mixed samples, two populations with distinct mean fluorescent intensities appeared, corresponding to the populations where either ligand binding occurred (SSTR2+) or was absent (SSTR2-). We analyzed the percentages of the two peaks and compared them to the percentage of SSTR2+ cells that were added to the mixed cultures. The experiment was repeated three times.

### Tumor cell staining with an SSTR2 antibody

To determine the SSTR2 expression of tumor cell lines, HCT116-WT, HCT116-SSTR2 and H69 cells were seeded into FACS tubes at a concentration of 1×10_ cells/mL in 1 mL of FACS buffer. The cells were incubated with 10 µL of a human SSTR2 APC-conjugated antibody (R&D systems, FAB4224A) for 60 minutes at rt. Following incubation, the samples were washed three times with 1 mL of FACS buffer. After washing, 100 µL of DAPI (0.1 µg/mL) was added to each sample as a live/dead stain prior to analysis by flow cytometry. AR42J cells could not be stained with this method since they are of rat origin.

### Isolation of bone marrow mononuclear cells (BMMCs)

Human BM samples were obtained from femoral heads of patients undergoing hip replacement surgery. Written informed consent in accordance with the Declaration of Helsinki was obtained from all patients according to protocols approved by the ethics committee of the Technische Universität München (TUM 339/21). BMMC were isolated after mincing the femoral head in PBS and following Ficoll density gradient centrifugation (Biocoll Separating Solution density 1.077g/ml, Bio-sell, BS.L 6115). Cells were frozen in 45% Iscove’s Modified Dulbecco’s Medium (IMDM) with GlutaMAX (Gibco, 31980048), 45% FBS (Gibco, 10270106) and 10% DMSO (Serva, 20385) and stored in liquid nitrogen until further use.

### Ligand binding to HSPC cells

To investigate the binding of MMC(S2Cy5)-TOC/JR11 to BM cells, we isolated BMMCs as described above, determined cell concentration by trypan blue counting, and seeded them into FACS tubes in 1 mL FACS buffer. They were incubated with 5 µL (0.25 µg) APC-eFluor780-CD34 (Thermo Scientific, 47-0349-41), PerCP-eFluor710-CD38 (Thermo Scientific, 46-0388-42), PE-Cy7-CD45RA (Thermo Scientific, 25-0458-42) and PE-CD90 (Thermo Scientific, 12-0900-81) for 30 min at rt. After that, they were incubated with 0.25 µM MMC(S2Cy5)-JR11/TOC for 30 min at rt. For blocking, 25 µM DOTA-JR11 was added 15 min before MMC(S2Cy5)-JR11/TOC. Single stained samples were used for compensation. Samples were washed with 1 mL FACS buffer three times and 100 µL DAPI (0.1 µg/mL) was added as live/dead stain prior to flow cytometry. We recorded as many events as possible (1.5-8 million events per sample). We analyzed the mean fluorescence intensity within each identifiable subpopulation (CD34+, CD34-, HSC, MPP, MLP and CD34^+^/CD38^-^). Cell populations were identified according to the relevant cell surface markers (**Figure 2a**). A total of n=6 samples were analyzed.

### SSTR2 expression of BMMC

To determine SSTR2 expression in BM cells, we followed the same protocol as above, but instead of the multimodal ligands, they were incubated with SSTR2-APC antibody (R&D Systems, FAB4224A) at a concentration of 10 µL/10_ cells for 30 min at rt. Single-stained samples were used for compensation. The samples were washed three times with 1 mL of FACS buffer, and 100 µL of DAPI (0.1 µg/mL) was added as a live/dead stain prior to flow cytometry analysis. A total of n=3 samples were analyzed.

### Determination of intracellular localization of multimodal ligands via Imaging Flow Cytometry

Imaging Flow Cytometry was performed using an Image Stream X Mark I platform (Amnis-Merck-millipore) in standard configuration, equipped with 405 and 488 nm lasers for excitation and a 785 nm laser for a scatter signal with standard filter sets. INSPIRE software (Amnis) was used for acquisition and IDEAS software (Amnis Seattle, WA, USA) for analysis. BMMC isolation, CD-marker staining and MMC(S2Cy5)-JR11/TOC staining were carried out as described above. Single stain compensation tubes, as well as an unstained tube, were prepared alongside test samples. Around 100,000 events per sample were acquired.

### Determination of cell proliferation and cell death after radioligand treatment

BMMCs were thawed in 90% IMDM with GlutaMAX, 10% FBS, and 20 µg/ml DNAse. Cells were washed and resuspended in cold FACS buffer for CD34+ isolation using the Miltenyi Biotec CD34 MicroBead Kit (#130-046-702) as per manufacturer instructions. CellTrace™ Violet (CTV, Thermo Scientific, C34557) staining was conducted based on the CD34^+^ cell count (0.05 µl CTV/ml/ 1.5×10^6^ cells) to determine the number of divisions undergone by a cell at the time of analysis. Following a 20 min incubation at 37°C, the cells were centrifuged and resuspended in serum-free medium composed of IMDM with GlutaMAX and 20% BIT9500 (Stem Cell Technologies, 09500) supplemented with three cytokines involved in stem cell maintenance: SCF (50 ng/ml, PeproTech, 300-07), TPO (12.5ng/ml, PeproTech, 300-18), andFlt3-L (50ng/ml, PeproTech, 300-19). 30,000 cells were seeded in 24-well plates and analyzed by flow cytometry either immediately (Day 0) or after a 4-day incubation period at 37°C with 5% CO2 (Day 4) following 30 min antibody staining and additional 15 min Annexin V APC binding (Biolegend, 640920). The antibodies used were as follow: CD34-APC-Cy7 (Biolegend, 343514), CD38-FITC (Biolegend, 356610), CD90-PE-Cy5 (Biolegend, 328112), CD123-PE (Biolegend, 306006), CD45RA-PE-Cy7 (Biolegend, 202316).

For radioligand treatment, cells were treated with 50 kBq/well (500 µl/well; 200 MBq/nmol) of ^177^Lu-DOTA-TOC or ^177^Lu-DOTA-JR11 for the entire incubation period of 4 days. Untreated cells served as controls. A total of n=16 samples were analyzed.

In this experiment, flow cytometry was conducted on a Miltenyi MACSQuant analyzer equipped with three lasers - blue (488 nm, 30 mW), red (640 nm, 20mW), and violet (405 nm, 40 mW). For each sample, 50,000 events were recorded. MACSQuantify™ Software was used for data acquisition and Flowjo 9.0-10.10 (BD Biosciences, USA) was employed for flow cytometric data analysis. The resulting data showed the distribution of cells (in percent) over up to 7 divisions per dataset. To simplify the comparison between groups, we calculated a “proliferation score” per sample. The proliferation score was defined as the weighted sum of all proliferations using the formula: “Proliferation score = 1*(% one division)+2*(% two divisions)+3*(% three division)+4*(% four division)+5*(% five divisions)+6*(% six division) +7* (% > six divisions). A higher weighted sum indicated more cell proliferation events.

### Statistical analysis

Data were plotted and statistically analyzed using GraphPad Prism 10 (GraphPad Software, US). Data are displayed as mean values and standard deviation if not otherwise specified. K_D_ values were obtained by applying a non-linear regression curve fit (Saturation binding – One site specific binding) in GraphPad Prism 10. We applied a paired t-test with correction for multiple comparisons to analyze multimodal ligand binding to HSPC subpopulations and cell proliferation and cell death data. Results with p-values p<0.05 were considered statistically significant. We performed linear regression analysis to correlate SSTR2-antibody binding to uptake of MMC(S2Cy5)-ligands.

## Acknowledgements.

We thank Jürgen Ruland and the Cell Analysis Core Facility at the Central Institute for Translational Cancer Research (TranslaTUM) at Technical University of Munich for instrument access and experimental support during flow cytometry. We thank Melpomeni Fani for providing a sample of DOTA-JR11 in the initial stage of the project. We extend our gratitude to Isotope Technologies Munich (ITM) for providing Lutetium-177 within our research collaboration.

## Author contribution statement

Conceptualization and design, A.A., W.A.W., K.S.G., S.K. Development of methodology, N.N., Y.M., J.R., M.B., A.A., W.A.W., K.S.G., S.K. Experiments and acquisition of data, N.N., Y.M., J.R., M.v.d.G., S.G., L.M.B., M.B., F.B., A.A., W.A.W., K.S.G. S.K. Analysis and interpretation of data, N.N., Y.M., J.R., A.A., W.A.W., F.B., K.S.G. S.K. Writing, review and/or revision of the paper, N.N., Y.M., J.R., M.v.d.G., S.G., L.M.B., M.B., F.B., A.A., W.A.W., K.S.G., S.K.

## Materials and correspondence

Correspondence and data or material requests should be addressed to

## Data availability

All data are available upon request to

