## Supplementary Figures and Tables for "Limitations of the radiotheranostic concept in neuroendocrine tumors due to lineage-dependent somatostatin receptor expression on hematopoietic stem and progenitor cells"

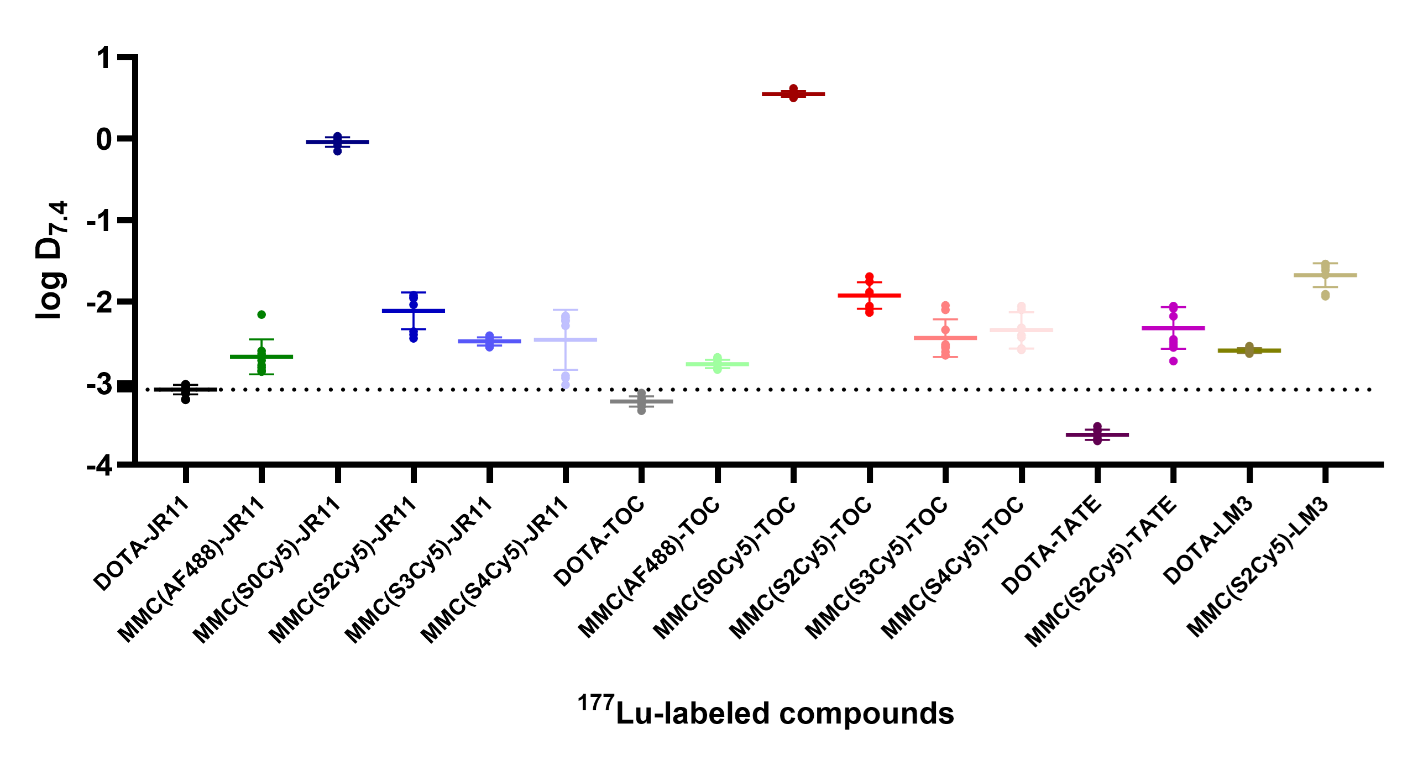

**Supplementary Figure 1**: Determination of the log D_7.4_ value (lower values = more hydrophilic) of the MMC(Dye)-ligands TOC, TATE, JR11 and LM3 and their respective DOTA-peptide.

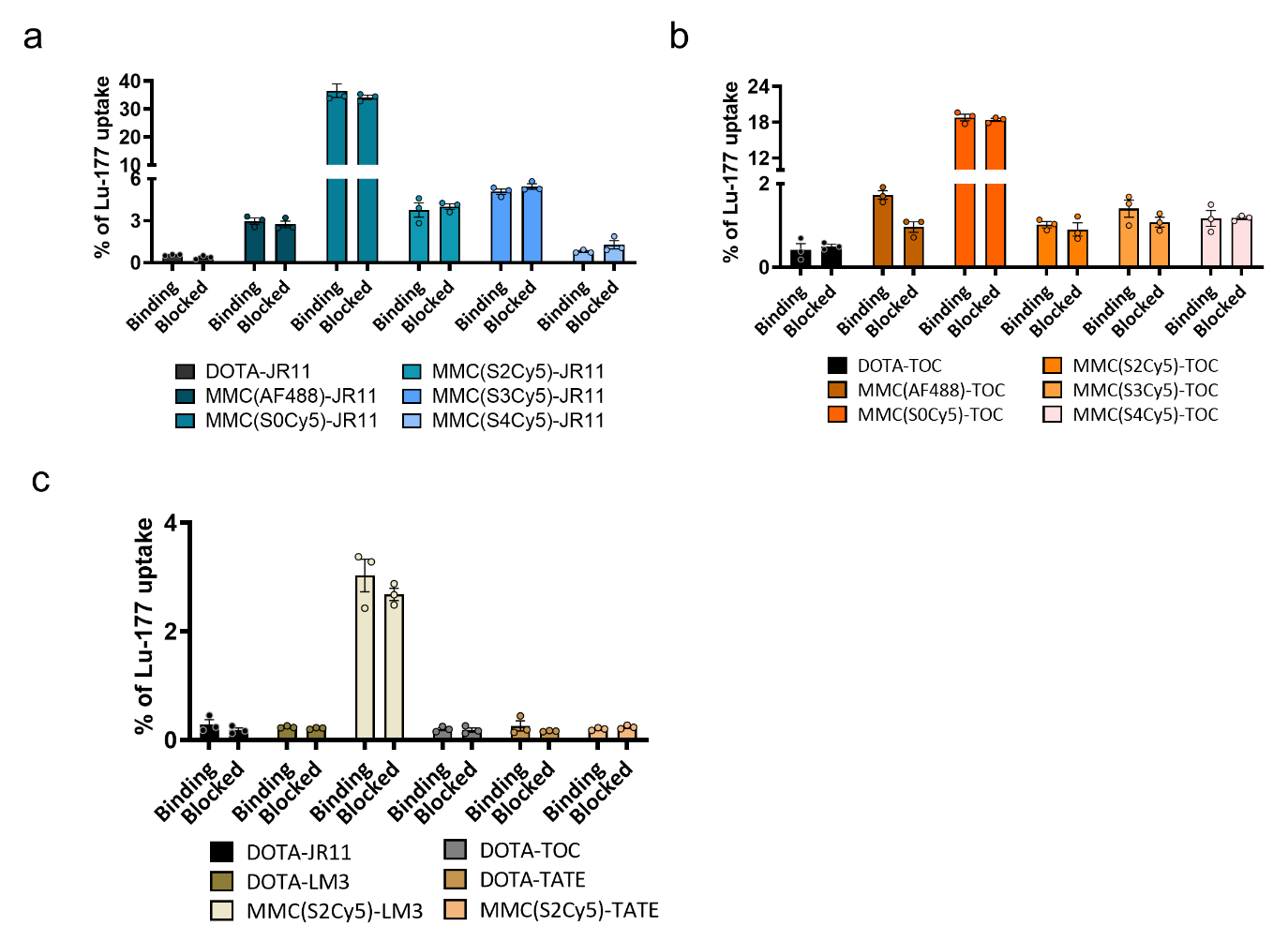

**Supplementary Figure 2:** Radioligand characterization on HCT116-WT cells. Cellular uptake of ^177^Lu-labeled JR11 **(a)**, TOC **(b),** LM3 and TATE **(c)** analogs, reported as % uptake of total added ^177^Lu-activity on SSTR2-negative HCT116-WT cells with or without blocking with a 1000-fold excess of DOTA-JR11.

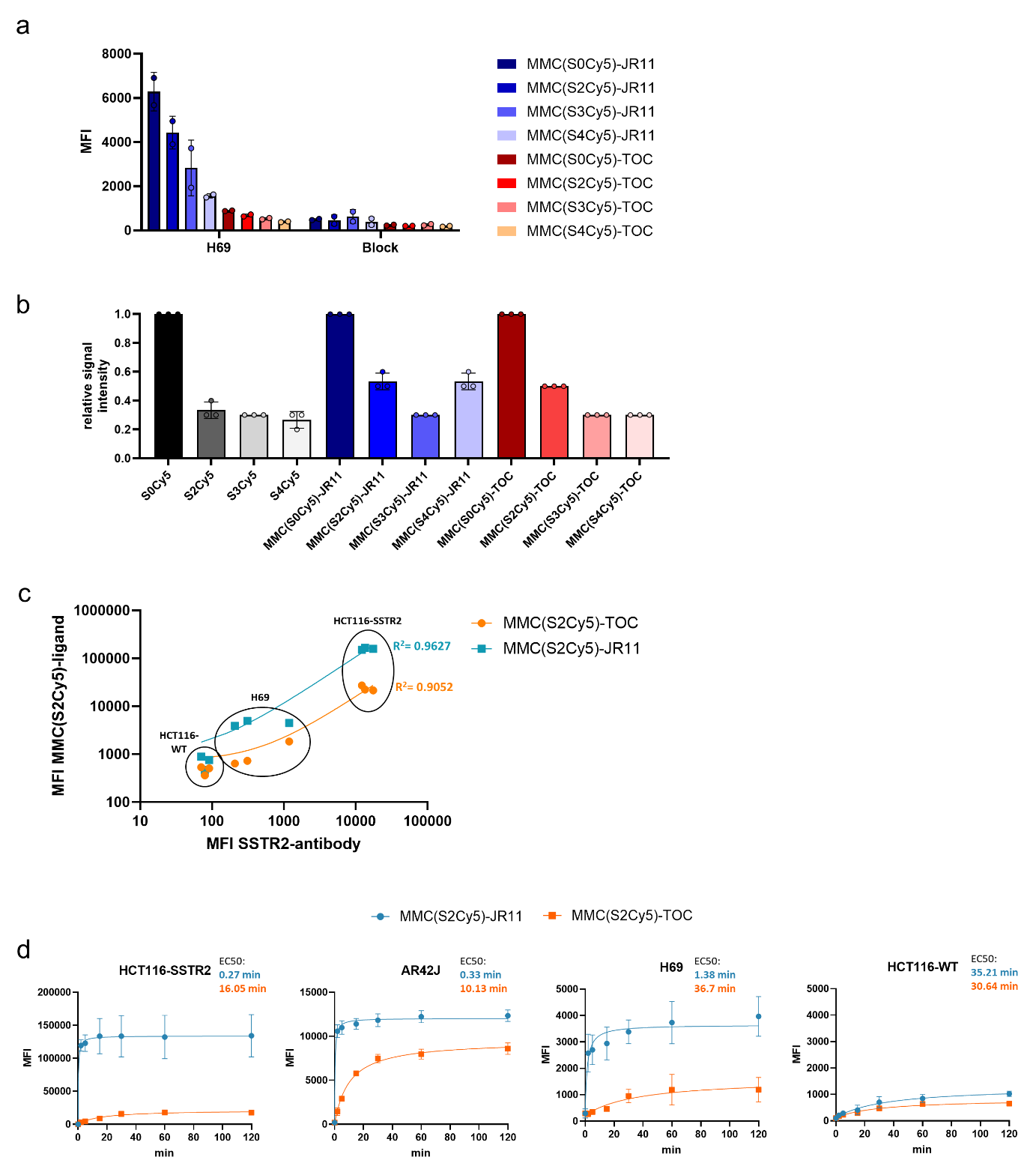

**Supplementary Figure 3**: **a)** Cellular uptake of MMC(Dye)-ligand reported as mean fluorescence intensity (MFI) on AR42J and H69 cells with or without blocking with 100-fold excess DOTA-JR11. **b)** Fluorescence intensity comparison of MMC(Dye)-conjugates and unconjugated DBCO-dyes in 5 µM DMSO. Intensities were normalized to the brightest dye (S0Cy5-DBCO). **c)** SSTR2 antibody binding in correlation to multimodal ligand binding in tumor cells. R^2^ values were derived from linear regression. **d)** Time-dependent cellular uptake of MMC(S2Cy5)-ligands reported as mean fluorescence intensity (MFI) for HCT116-SSTR2, AR42-J, H69 and negative control cell line HCT116-WT. JR11-based compounds show higher MFI and faster cell association in SSTR2-expressing cell lines as determined by EC_50_ values.

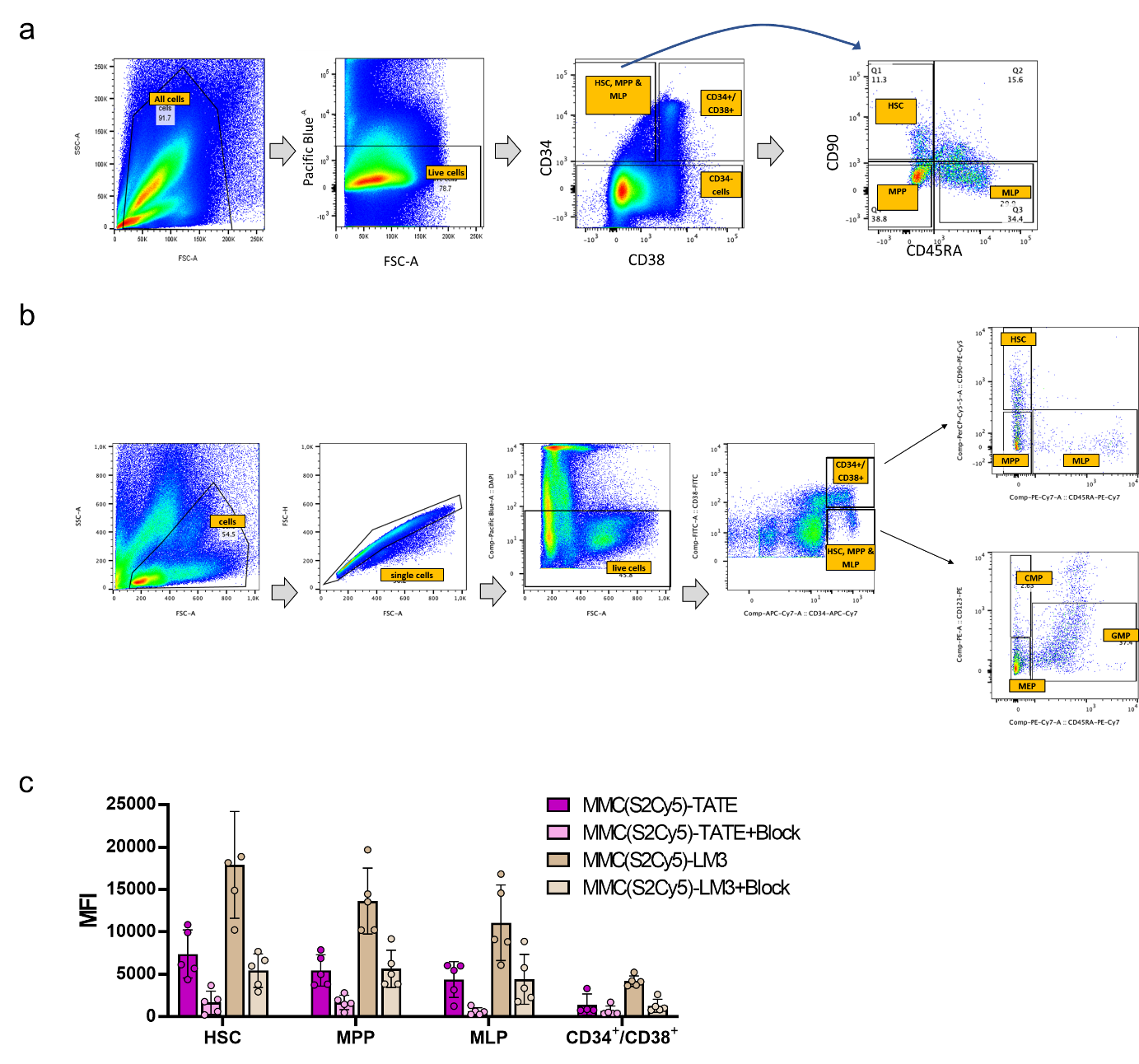

**Supplementary Figure 4:** a) Multicolor flow cytometry gating strategy were employed for the identification of the 4 human HSC subpopulations HSC, MPP, MLP and CD34+/CD38+ based on CD-markers CD34, CD38, CD90 and CD45RA. b) Multicolor flow cytometry gating strategy for the identification of the 6 human HSC subpopulations HSC, MPP, MLP and CMP, MEP and GMP (together they represent CD34+/CD38+) based on CD-markers CD34, CD38, CD90, CD45RA and CD123. c) MFI of MMC(S2Cy5)-TATE/LM3 in HSPC subpopulations with or without blocking.

**Supplementary Table:**

Supplementary Table 1: Age and gender distribution of the treatment cohort. Age in years (y) is given as mean and (range).

|  | Total | Female | Male |
| --- | --- | --- | --- |
| Number (n) | 16 | 8 | 8 |
| Age (y) | 52,875 (32-59) | 50,375 (32-59) | 55,375 (46-58) |
| High Sensitivity (n) | 8 | 4 | 4 |
| Low Sensitivity cohort (n) | 8 | 4 | 4 |
| High Sensitivity (y) | 50,25 | 47,0 (32-55) | 53,5 (46-58) |
| Low Sensitivity cohort (y) | 55,5 | 53,75 (46-58) | 57,25 (55-58) |
